# Packaging of supplemented urokinase into naked alpha-granules of *in vitro*-grown megakaryocytes for targeted therapeutic delivery

**DOI:** 10.1101/2023.12.05.570278

**Authors:** Mortimer Poncz, Sergei V. Zaitsev, Hyunsook Ahn, M. Anna Kowalska, Khalil Bdeir, Rodney M. Camire, Douglas B. Cines, Victoria Stepanova

## Abstract

Our prior finding that uPA endogenously expressed and stored in the platelets of transgenic mice prevented thrombus formation without causing bleeding, prompted us to develop a potentially clinically relevant means of generating anti-thrombotic human platelets *in vitro* from CD34^+^ hematopoietic cell-derived megakaryocytes. CD34^+^-megakaryocytes internalize and store in α-granules single-chain uPA (scuPA) and a uPA variant modified to be plasmin-resistant, but thrombin-activatable, (uPAT). Both uPAs co-localized with internalized factor V (FV), fibrinogen and plasminogen, low-density lipoprotein receptor-related protein 1 (LRP1), and interferon-induced transmembrane protein 3 (IFITM3), but not with endogenous von Willebrand factor (VWF). Endocytosis of uPA by CD34^+^-\megakaryocytes was mediated in part via LRP1 and αIIbβ3. scuPA-containing megakaryocytes degraded endocytosed intragranular FV, but not endogenous VWF, in the presence of internalized plasminogen, whereas uPAT-megakaryocytes did not significantly degrade either protein. We used a carotid-artery injury model in NOD-scid IL2rγnull (NSG) mice homozygous for VWF^R1326H^ (a mutation switching binding VWF specificity from mouse to human glycoprotein IbmlIX) to test whether platelets derived from scuPA-MKs or uPAT-Mks would prevent thrombus formation. NSG/VWF^R1326H^ mice exhibited a lower thrombotic burden after carotid artery injury compared to NSG mice unless infused with human platelets or MKs, whereas intravenous injection of either uPA-containing megakaryocytes into NSG/VWF^R1326H^ generated sufficient uPA-containing human platelets to lyse nascent thrombi. These studies suggest the potential to deliver uPA or potentially other ectopic proteins within platelet α-granules from *in vitro-*generated megakaryocytes.

**Key points:**

- Unlike platelets, in vitro-grown megakaryocytes can store exogenous uPA in its α-granules.
- uPA uptake involves LRP1 and αIIbβ3 receptors and is functionally available from activated platelets.

## Introduction

There is an unmet need for safe and effective therapeutics to treat or prevent thromboembolism in settings such as post-surgery settings, extensive trauma or recent bleeds. Yet, the risk of perioperative bleeding often delays the introduction of anticoagulation during this critical period. Similar tradeoffs in the timing of anticoagulation based on the risk of bleeding arise in patients with ischemic stroke, after traumatic brain injury and extensive trauma. In these and in related settings, there is a need for therapies that distinguish preexisting hemostatic clots, leaving them intact, while targeting for fibrinolysis nascent, potentially occlusive, thrombi.

Fibrinolytic agents, such as the plasminogen activators (PA) tissue-type PA (tPA) and urokinase PA (uPA) are effective at lysing thrombi^1–3^, but permeate preexisting hemostatic fibrin^4^ and can degrade circulating fibrinogen^5^. Moreover, both PAs have short half-lives, necessitating continuous intravenous infusions for prophylaxis^6^ and have vasoactive and neurotoxic side effects due to signaling through identified cell-surface receptors and diffusion through an injured blood-brain barrier such as after a stroke (reviewed in ^7^). Therefore, the clinical use of fibrinolytic agents has been limited primarily to the management of a subset of patients with acute ischemic stroke or acute life- or limb-threatening thromboembolism^8–10^.

To help rectify these limitations, we found that PAs linked to red blood cells (RBCs) or platelets provides protracted and safe thromboprophylaxis in several animal models^11,12^. However, RBCs are not rapidly and selectively recruited to sites of incipient arterial thrombus formation^13^, and linking the PAs to the surface of the platelets can lead to thrombocytopenia and bleeding^11^.

As an alternative approach, we expressed single-chain (sc) murine uPA endogenously in platelets of transgenic mice. The scuPA was stored in platelet α-granules without causing systemic fibrinolysis. The transgenic mice had a mild bleeding diathesis during parturition^14^ ameliorated by tranexamic acid (TEA), which inhibits plasmin and uPA activity^15^. These mice simulated human Quebec Platelet Disorder (QPD), which is caused by a duplication of the uPA (*PLAU*) gene, also characterized by storage of uPA in platelet α-granules^16–19^, absence of systemic fibrinolysis, and a mild bleeding diathesis mitigated by TEA^14^. We have previously shown that infusion of the uPA-containing platelets (uPA-PLTs) into wildtype mice prevented nascent thrombus growth in a microemboli infusion model^14^. This raises the possibility that delivering uPA packaged within the α granules of platelets, which naturally target sites of incipient clot formation, inverts their function from prothrombotic to fibrinolytic and does so in a manner that does not lyse mature clots that are no longer recruiting platelets^14^.

Recent advances in the generation of platelets from *in vitro*-grown megakaryocytes have been made, especially beginning with induced pluripotent stem cells (iPSCs)^20–22^. Indeed, a recent paper describing the infusion of platelets generated from iPSCs into a patient as a platelet transfusion had been described^23^. We wondered if *in vitro*-grown megakaryocytes can be used to deliver uPA as the megakaryocytes are grown in defined medium and thus lack the normal array of endocytosed proteins. Prior studies have shown that these in vitro -grown megakaryocytes can endocytose added fibrinogen and factor (F) V^23^. We speculated that these megakaryocytes may have “naked α-granules” and be especially able to endocytose other proteins, and thus may be useful for modifying before being given to a patient as a novel therapeutic. Here, we show that scuPA or a thrombin-, but not plasmin-activatable, uPA (uPAT)^11^ added to the media were readily endocytosed by maturing megakaryocytes. We define receptors involved in their uptake and the interaction of these uPAs with other endocytosed and with endogenous α-granular proteins. Using an *in vivo* murine carotid artery photochemical injury model, we show that the uPAs studied can effectively prevent thrombus development. We discuss potential uses of this approach for targeted thromboprophylactic fibrinolysis and the application of this strategy for delivering other ectopic therapeutics via platelets.

## Materials and methods

### Supplemental methods

Please see the Supplement for a description of reagents and antibodies.

### Generation of megakaryocytes

Granulocyte colony-stimulating factor–mobilized human CD34^+^-hematopoietic progenitor cells (HPCs), purchased from the Fred Hutchinson Cancer Research Center Hematopoietic Cell Processing and Repository Core, were differentiated into megakaryocytes (CD34^+^-MKs) in medium composed of 80% Iscove-modified Dulbecco medium, 20% bovine serum albumin/insulin/transferrin serum substitute, 20 μg/mL low-density lipoproteins, 100 μM 2-mercaptoethanol with the following recombinant human cytokines added: 1 ng/mL stem cell factor, 100 ng/mL thrombopoietin, 13.5 ng/mL interleukin-9 and 7.5 ng/mL interleukin 6, as described by us^24^. CD34^+^-HPCs, differentiated *in vitro* into megakaryocytes, were characterized as in ^25^.

### Generation of uPA-containing megakaryocytes (uPA-MKs) and fluoro-immunocytochemical studies

Day (d) 10 or d11 CD34^+^-derived megakaryocytes (CD34^+^-MKs) were incubated for various times in complete growth medium containing various combinations of the following fluorophore-conjugated proteins: Alexa-488 or Alexa-555 scuPA, Alexa-488 uPAT, Alexa-647 noncleavable plasminogen (ncPLG)^26^, and Alexa-568 Factor (F) V. The CD34^+^-MKs were washed once with phosphate-buffered saline (PBS) and immobilized on microscopic slides using Shandon CytoSpin III Cytocentrifuge with the M964-20FW-CytoSep™ Double Funnel for 15 minutes. To stain endogenous or endocytosed unlabeled proteins, megakaryocytes, immobilized on slides after the cytospin were fixed in 4% paraformaldehyde (PFA) for 15 minutes, permeabilized in PBS/0.1% Triton X-100 (Tx-100), and 1% bovine serum albumin (BSA) blocking solution was added for 1 hour. von Willebrand Factor (VWF) and/or human uPA were detected using rabbit anti-VWF or anti-IFITM-3 polyclonal or mouse anti-uPA monoclonal antibodies, respectively, rabbit and mouse control antibody

Megakaryocytes were examined using a confocal laser-scanning microscope (Zeiss LSM 710; Carl Zeiss) equipped with a Plan Apo 40× water-immersion objective lens (NA 1.2). The Z-stack distance between the slices was set as 0.3 µm, with a 1024 × 1024 pixel resolution for each slice. Three-dimensional reconstruction and maximal projection were performed using Volocity 6.3 software (Perkin Elmer). In most experiments, the Manders overlaps coefficients were used to determine the proportion of signals from scuPA that coincided with signals from uPAT, or signals from scuPA or uPAT that coincided with signals from target markers. Values close to 1 indicate that a large proportion of uPAs signals overlapped with signals from the target markers^27^.

### Flow cytometry studies of protein uptake

CD34^+^-MKs differentiated for 10-11 days were incubated in complete growth medium containing a combination of the following fluorophore-conjugated proteins: Alexa-488 or Alexa-568 scuPA, Alexa-488 uPAT, Alexa-647 ncPLG, Alexa-568 FV and Alexa-555 receptor-associated protein linked to human IgG Fc fragment (FcRAP)^28^. Unlabelled, recombinant FcRAP (1 μM) or the blocking αIIbβ3 monoclonal antibody ReoPro (Abciximab) (200 µg/mL), added 1 h prior of scuPA, uPAT and FV to the cells and incubated for 8 and 24 h, were used as an LRP1 and αIIbβ3 antagonists, respectively, to compete with scuPA, uPAT and FV uptake. The cells were washed with PBS, stained with allophycocyanin (APC)-conjugated anti-CD42b to gate the matured MKs, and the uptake of endocytosed fluorescent proteins was analyzed in CD42b-positive population using a CytoFlex S flow Cytometer (Beckman).

### Western blotting of α-granule proteins

CD34^+^-MKs differentiated for 10 days were incubated with purified plasmin-free plasminogen at 20 µg/mL for 24 hours, washed and then incubated for an additional 18 hours with recombinant human scuPA or uPAT (200 nM each)^11^ in the absence or presence of human FV (400 nM)^29^. The cells were then lysed in 1×radioimmunoprecipitation assay (RIPA) buffer with added 1× proteinase inhibitor mixture (Sigma) and 1× phosphatase inhibitors mixture 2 (Sigma). Protein concentrations in cell lysates were measured using the Bradford Protein Assay. Lysates were subjected to SDS-PAGE and Western blotting using the following antibodies: rabbit anti-human uPA polyclonal antibodies; rabbit anti-human VWF polyclonal antibodies; mouse anti-human FV monoclonal antibody; and HRP-conjugated anti-β-actin rabbit polyclonal antibody. Secondary horse radish peroxidase-conjugated goat anti-mouse and anti-rabbit polyclonal antibodieswere used for detection, and protein bands were visualized using SuperSignal™ West Femto Maximum Sensitivity Substrate. Luminescent signals on the blots were scanned and quantified using a Li-COR C-Digit Blot scanner (LI-COR Biosciences)/Image Studio™ software (LI-COR Biosciences).

### In vivo-murine studies

NOD-scid IL2rγnull (NSG) mice were originally purchased from Jackson Laboratory. The transgenic NSG/VWF^R1326H^ mice, homozygous for a CRISPR/Cas9 mutation in murine VWF, was previously described^25,30^. Carotid artery thrombosis was induced by photochemical injury, as described by us^31^. Briefly, adult male and female mice were anesthetized with Phenobarbital (80 mg/kg), and the right carotid artery and left jugular vein were exposed by blunt dissection. A 3 mW, 540-nm laser beam (green) was applied to the artery from a distance of 5 cm. Rose bengal dye (50 mg/kg body weight) was injected into the left jugular vein and blood flow in the artery was recorded using a small animal blood flow meter (model T106; Transonic Systems) over the ensuing 40 minutes. Time to initial formation of a stable completely occlusive thrombus (occlusion > 10 minutes) was used as the endpoint. Four hours before injecting rose bengal dye, some animals were injected intravenously with 3×10^6^ CD34^+^-MKs in 100 μL PBS. Prior to infusion, MKs were or were not pre-loaded with recombinant scuPA or uPAT as follows: day 10 CD34+ MKs were left intact or incubated with recombinant scuPA or uPAT (600 nM each) for 24h in the culture media. The cells were washed, and re-suspended in PBS at 3×10^7^ CD34^+^-MKs/ml concentration.. All murine studies were done after institutional Animal Care and Use Committee. Euthanasia was performed as approved by the Panel on Euthanasia of the American Veterinary Medical Association.

### Statistical analysis

Means ± standard error of the mean (SEM) are shown. For data obtained by flow cytometry differences were analyzed by ordinary one-way Anova. For *in vivo* studies, differences in carotid blood flow in the photochemical carotid injury studies were analyzed by ordinary one-way analysis of variance (ANOVA) using GraphPad Prism 5. For all studies, differences were considered significant if the *P* value was <0.05 compared to the indicated control.

### Institutional approval of studies

Approval was obtained from the institutional internal review board for the volunteer blood samples and from the animal care and use committee for murine studies. The study was conducted in accordance with the Declaration of Helsinki.

## Results

### Endocytosis of uPA by in-vitro-grown megakaryocytes

We and others have shown that *in vitro*-generated MKs differentiated in media can endocytose fibrinogen, FVIII and FV until the late stages of differentiation^23,32,33^; however, given the lack of these proteins in the media when megakaryocytes are grown, these proteins would be absent from the α-granules. Perhaps if uPA were added to the media that they would endocytose the uPA instead and potential be useful to generate uPA-PLTs for targeted thrombolysis. We, therefore, added to the media scuPA. This uPA contains the N-terminal uPA-receptor (uPAR)-binding growth factor-like domain (GFD), a kringle domain (KD) and a C-terminal pro-protease domain (Figure 1A, top). scuPA is dependent on uPAR as well as low-density lipoprotein receptor-related protein 1 (LRP1) for uptake in cells^34^. Separately, we also added a low-molecular weight variant we developed^11^, designated uPAT, which lacks the two N-terminal domains (Figure 1A lower left panel), which makes it incapable of binding to uPAR. uPAT is activated by thrombin, e.g., generated at the site of clot formation, but not by plasmin^34^, and this may prevent uPAT activation by plasmin, avoiding α-granular protein degradation seen in QPD^35^.

**Figure 1.**
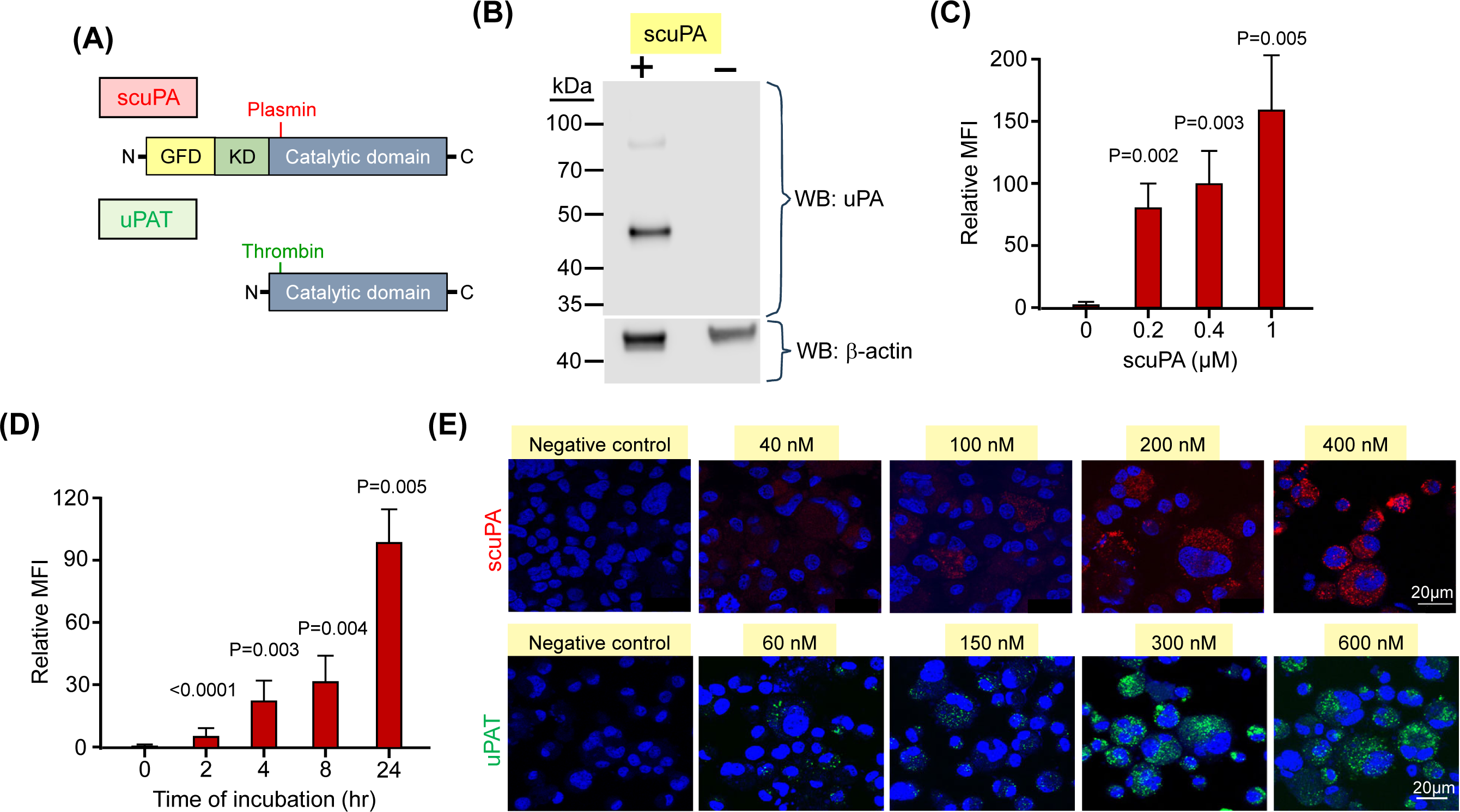
uPA is endocytosed by *in vitro*-grown Mks and stored in granules. (**A**) Schematics of the structures of scuPA (top) and uPAT (bottom). Full-length scuPA is composed of *i.* an N-terminal growth factor-like domain (GFD, yellow) that binds uPAR; *ii.* a unique kringle domain (K, green) that mediates LRP1-dependent intracellular uptake^59^, binding to integrins^60,61^ and nuclear translocation^59^ and *iii.* the protease domain (Catalytic domain, grey)^62^. Site of activation by plasmin is shown in red. uPAT is composed of the protease domain in which ^157^F^158^K has been deleted^51^ to create a thrombin cleavage/activation site (shown in green) (**B**) Representative western blot (WB) of lysates of *in vitro*-grown d11 megakaryocytes in the presence (+) or absence (-) of scuPA (400 nM) added at d10. The WB at the top was with anti-human uPA mouse monoclonal antibody followed by HRP-conjugated goat anti-mouse antibody and bottom shows HPR-conjugated anti-β-actin antibody as a loading control. Size marker is shown to the left of the blot. (**C**) Dose-dependent uptake of Alexa-568 scuPA by CD34^+^-MKs. Y axes denote mean fluorescence intensity (MFI) measured by flow cytometry. Mean ± 1 standard deviation (SD) of relative uptake of scuPA compared to the values in its absence. N = 4 independent studies. P values were determined by ordinary one-way ANOVA. (**D**) Same as (C), but for time course of Alexa-568 scuPA (400 nM) uptake by the CD34^+^-Mks. **(E)** Visualization using confocal microscopy of Alexa-568 scuPA (top, red) or Alexa-488 uPAT (bottom, green) endocytosed by d11 CD34^+^-Mks for 24 hours starting on d10 initiation of culture. DAPI (blue) depicts the nuclei. Scale bar is shown.

Western blotting of cell lysates showed that control MK were devoid of uPA, whereas cells incubated with scuPA showed the presence of the taken up protein (Figure 1B). Both Alexa-568-conjugated scuPA and Alexa-488-conjugated uPAT were internalized by d10 CD34^+^-MKs in a concentration-dependent (Figures 1C and Supplement (S) 1A, respectively) and in a time-dependent (Figure 1D and S1B, respectively) manner. Both appeared as punctate particles in the MKs (Figure 1E). In contrast, isolated human platelets did not take up either uPAs (Figure S2).

### Pathways of uPA uptake by in vitro-generated megakaryocytes

One can suggest that scuPA and uPAT, based on their structural differences, enter megakaryocytes through different pathways and are stored in distinct granules. Alexa-568 scuPA and Alexa-488 uPAT were co-incubated with CD34^+^-MKs for 24 hours. Both scuPA and uPAT resided in the same granules as visualized by confocal microscopy with representative data shown in Figure 2A and Video 1.

**Figure 2.**
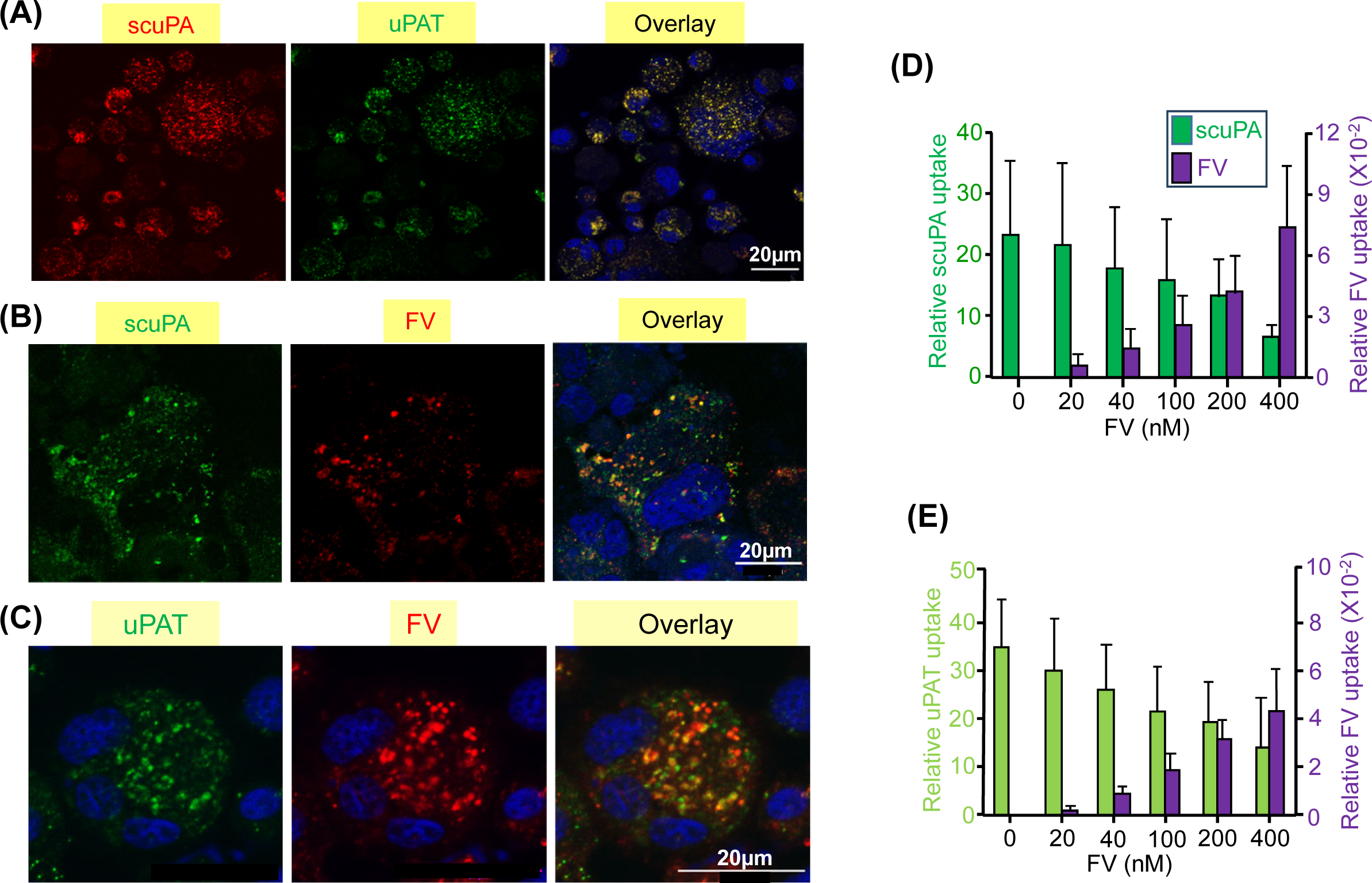
scuPA, uPAT and FV share an endocytic pathway in CD34^+^-MKs. **(A)** Representative confocal images of d10-CD34^+^-MKs loaded simultaneously by preincubation with Alexa-568 scuPA (red) and Alexa-488 uPA-T (green) for 24 hours. Nuclear staining by DAPI is depicted by blue. Overlap is in yellow and is shown to the right. Scale bar is shown. Quantitative analysis of overlap is shown in Table 1. (**B** and **C**) Similar confocal image studies as in (A) but of CD34^+^ MKs preincubated with Alexa-488 scuPA (B) or Alexa-488 uPAT (C) for 24 hours and then incubated with Alexa-568 FV for 2 hours (red) on d11. (**D** and **E**) Uptake of Alexa-488 scuPA (D) or Alexa-488 uPAT (E), each at 400 nM, by d11-CD34^+^-MKs in the absence or presence of Alexa-568-FV (0-400 nM). Y axis denotes MFI measured by flow cytometry. Mean ± 1 SD are shown. N = 4 independent experiments.

### Exogenous uPA is stored in α-granules with other proteins added to the media

FV is endocytosed by human MKs and stored in α-granules^36^. To help elucidate whether uPAs endocytosed by MKs are also stored in α-granules, CD34^+^-MKs were incubated with exogenously added Alexa-488 scuPA or uPAT for 18 hours, followed by co-incubation with Alexa-568 FV for an additional 2 hours. By confocal microscopy, there was extensive, but not as complete overlap as seen in Figure 2A, indicating that endocytosed scuPA or uPAT are stored in many of the same α-granules as endocytosed FV (Figures 2B and 2C, respectively, and Video 2). Furthermore, added Alexa-568 FV competed for the uptake of Alexa-488 scuPA and uPAT in the *in vitro*-grown CD34^+^-MKs in a concentration-dependent manner (Figures 2D and 2E, respectively), further supporting that the uPAs are stored in α-granules that contain physiologically endocytosed proteins.

### uPA is taken up via LRP1

The finding that uPAT localizes within the same α-granules as scuPA and extensively co-localizes with FV led us to investigate the involvement of LRP1, which mediates the endocytosis of FV by CD34^+^-MKs^36^. We confirmed prior studies that LRP1 is expressed during late megakaryopoiesis but is lost prior to platelet release^37^. We show by Western blot that d10 CD34^+^-MKs, but not donor-derived platelets express LRP1 (Figure 3A). Internalized scuPA is co-localized with LRP-1 in d11 CD34^+^-MKs (Figure 3B). Internalized Alexa-488 scuPA co-localized with the internalized the LRP chaperone/antagonist FcRAP^28^ (Figure 3C), and partially inhibited the uptake by CD34^+^-MKs of Alexa-488 scuPA, uPAT and FV (Figure 3D).

**Figure 3.**
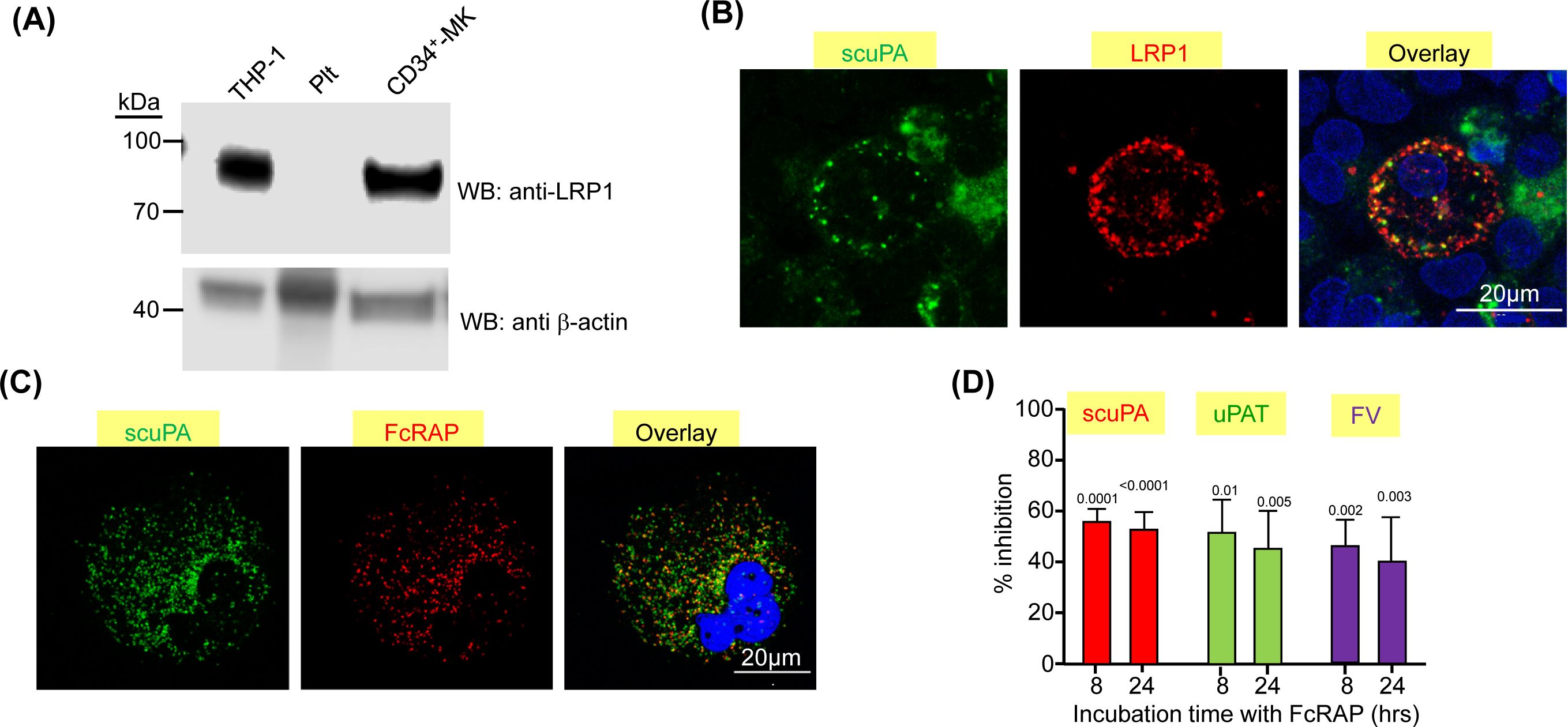
LRP1 mediates uptake of scuPA, uPA-T and FV by CD34+ MKs. **(A)** Representative WB, done as in Figure 1B, of lysates from HP-1 cell line as the positive control^63^, donor-derived platelets and d11-CD34^+^-MKs using anti-LRP light chain rabbit polyclonal antibody followed by HRP-conjugated goat anti-mouse antibody and HPR-conjugated anti-β-actin antibody as the loading control. (**B**) Confocal images of d11 CD34^+^-Mks preloaded with recombinant scuPA (1 μM) for 24 hours and stained with mouse anti-uPA monoclonal antibody and anti-LRP1 rabbit polyclonal antibody (Novus Biological) followed by Alexa-488 goat anti-mouse polyclonal antibody (green) and Alexa-555 goat anti-rabbit polyclonal antibody (red) and DAPI nuclear stain (blue). **(C)** Confocal images of d11-CD34^+^-MKs preloaded with Alexa 488 scuPA for 24 hours (green) followed by incubation with Alexa-555 FcRAP for 2 hours (red) on d11. Scale Bar = 20 μm. **(D)** Inhibition of scuPA, uPA-T and FV uptake by FcRAP. D11-CD34^+^-Mks were incubated with Alexa-488 scuPA or Alexa-488 uPAT or Alexa-568 FV (200 nM) in the absence or presence of FcRAP (2 µM) for either 8 or 24 hours and MFI was measured using flow cytometry. Y axis denotes % of uptake in the presence of FcRAP to in its absence. Mean ± 1 SD is shown. N = 3-4 independent studies. Data were analyzed using an ordinary one-way ANOVA.

### uPA and fibrinogen share an endocytic pathway in CD34^+^-MKs

On confocal microscopy, CD34^+^-MK co-incubated with Alexa-568 scuPA and Alexa-488 fibrinogen showed extensive co-localization of the proteins (Figure 4A, Video 3). Fibrinogen is known to be endocytosed via integrin αIIbβ3 in platelets and MKs^32,38^. The αIIbβ3-blocking antibody ReoPro^39,40^ reduced uptake of both fibrinogen and scuPA, but did not significantly affect endocytosis of FV (Figure 4B). Co-incubation with FcRAP and ReoPro did not further reduce uptake of Alexa-488 scuPA than FcRAP alone (Figure 4C), suggesting a common point in the pathway of uptake blocked by FcRAP and Abciximab. In support of this shared pathway, we observed extensive co-localization of the endocytosed scuPA both with LRP1 which carries out endocytosis of its ligands via the clathrin-coated pits^41^ and with the IFN-induced transmembrane protein 3 (IFITM3) (Figure 4D, Video 4), which mediates endocytosis of fibrinogen in CD34^+^-MKs and platelets through clathrin-coated pits via binding to clathrin and αIIbβ3^42^.

**Figure 4.**
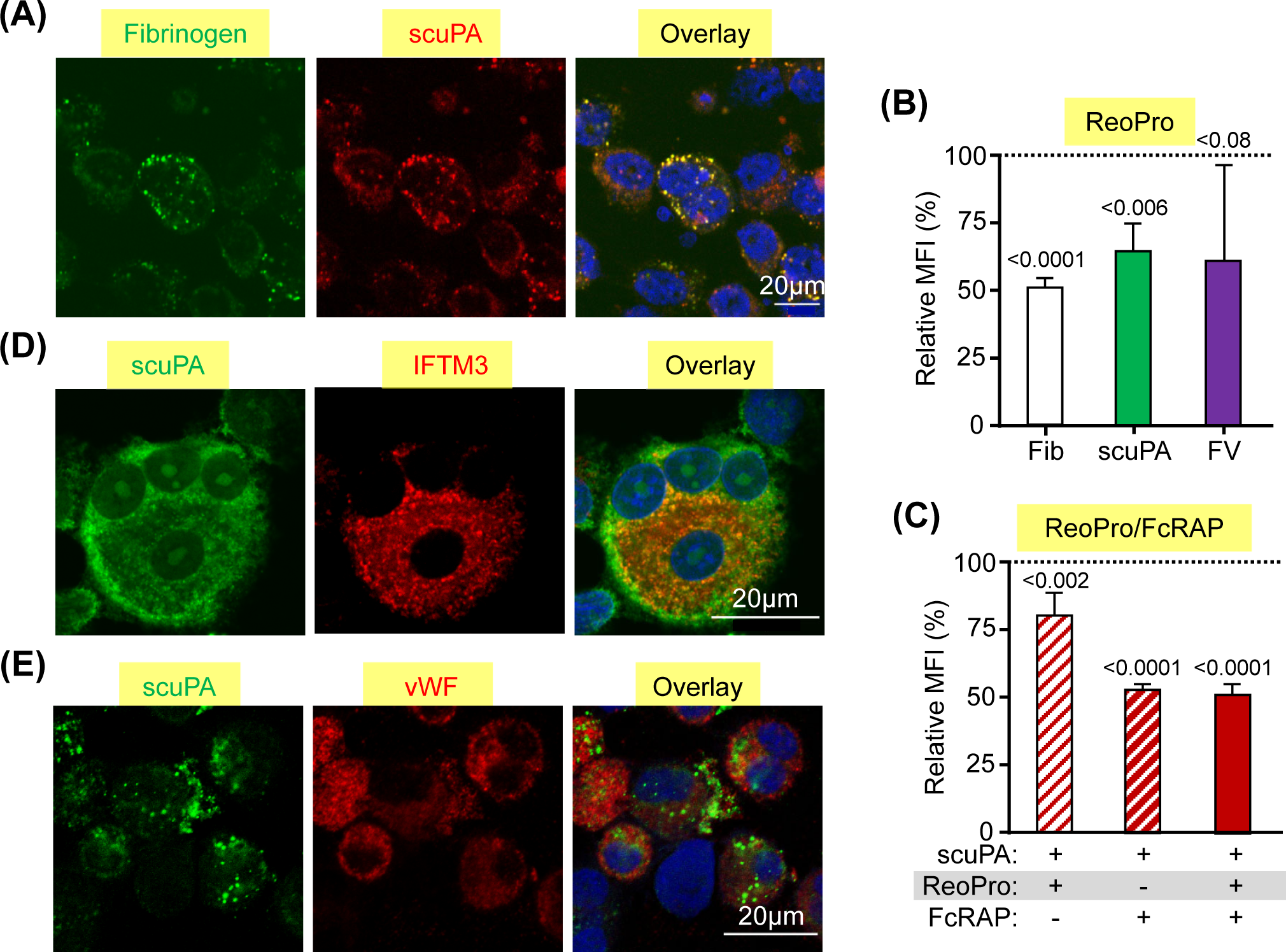
αIIbβ3 mediates uptake of scuPA by CD34^+^-MKs. **(A)** Confocal images of d11-CD34^+^-Mks loaded with Alexa-488 fibrinogen and Alexa-586 scuPA for 24 hours. Co-localization of scuPA and fibrinogen is depicted in yellow in the overlay panel (right). DAPI (blue) depicts the nuclei. Scale bar is shown. For quantitation of co-localization, see also Table 1. (**B** and **C**) In (B) Inhibition of fibrinogen, scuPA and FV (200 nM each) uptake by ReoPro monoclonal antibody (200 μg/mL). (C) is a similar study but of scuPA alone and inhibition was by ReoPro (200 µg/mL) and/or FcRAP (200 µg/mL). In both, Y axis denotes percent of uptake inhibition by ReoPro and/or FcRAP. Data were analyzed using the 2-way Anova test. Mean ± 1 SD is shown. N = 3-4 independent studies. Data were analyzed using a 2-way Student t-test. (**D** and **E**) Confocal studies of MKs loaded with scuPA (600 nM) and stained with Alexa 488 - mouse anti-uPA monoclonal antibody, and in (D), anti-IFITM3 rabbit polyclonal antibody or in (E), anti-VWF antibody mouse polyclonal antibody followed by Alexa-555 or Alexa-647 goat anti-rabbit polyclonal antibody (red) in (D). Nuclei were stained blue by DAPI. For quantitation of overlap, see also Table 1.

### Lack of co-localization of uPA and endogenously-expressed VWF

We next asked whether scuPA and/or uPAT endocytosed by CD34^+^-MKs co-localized with an endogenously-expressed protein stored in α-granules such as VWF^43^. We found very limited co-localization of endocytosed scuPA and uPAT with VWF (Figure 4E and S3, Video 5), suggesting that endocytosed uPAs and endogenously-expressed VWF are nearly completely segregated into different α-granules in MKs.

### Endocytosed scuPA and uPAT differ in degradation of α-granular proteins

In QPD and in platelet uPA-transgenic mice, α-granular proteins are proteolyzed by plasmin activated by the stored uPA, potentially compromising hemostasis^14,16–19^. Therefore, we investigated whether exogenously added uPAs also trigger proteolysis of proteins stored in the α-granules of *in vitro*-grown CD34^+^-Mks. As we did not detect plasminogen in the lysates of *in vitro*-grown CD34^+^-MKs (Figure S4), we inferred that these cells would only contain plasminogen *in vivo* if it were endocytosed from plasma. To begin to study the effect of plasminogen on uPA-loaded Mks, we used CD34^+^-MKs pre-loaded with Alexa-488 scuPA and Alexa-568 FV for 18 hours and then added Alexa-647-conjugated ncPLG, which cannot be converted to active plasmin^26^, for 18 hours. The use of ncPLG was to avoid activation/degradation of endocytosed scuPA and FV with subsequent trafficking to a degradation rather than a storage pathway. We found significant co-localization of endocytosed scuPA, FV and ncPLG in the α-granules of CD34^+^-MKs (Figure 5A).

**Figure 5.**
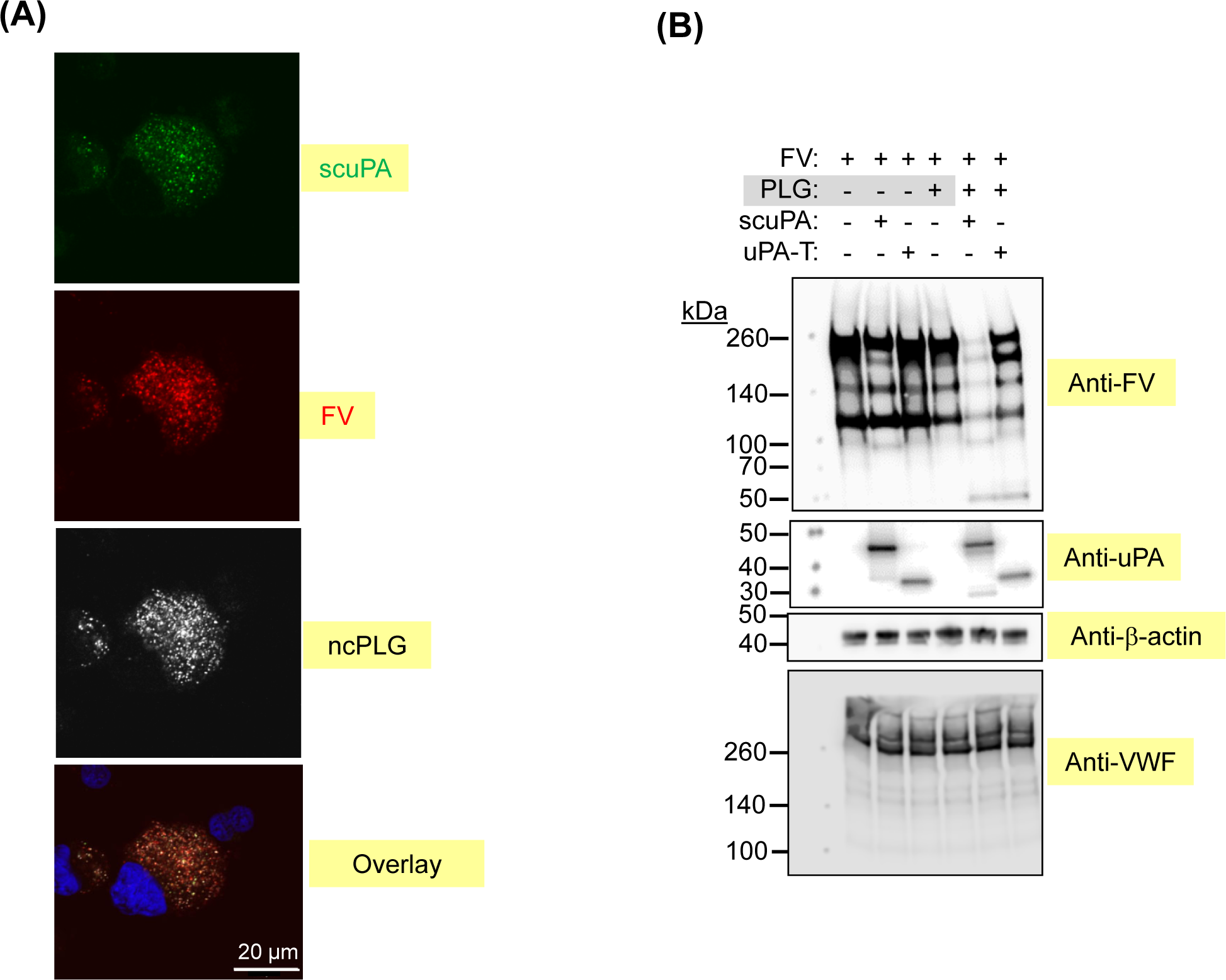
Effects of PLG on uPA-MKs. (**A**) Confocal images of d11-CD34^+^-Mks preincubated for 24 hours with Alexa-488 scuPA (green), Alexa-568 FV (red) and Alexa-647 ncPLG (white). Nuclei were stained blue by DAPI. For quantitation of overlap, see also Table 1. (**B** and **C**) WB analysis of lysates prepared from d11-CD34^+^-Mks with or without loading with enzymatically active PLG (20 µg/mL) on d10 for 18 hours followed by a wash and incubation with scuPA (B) or uPAT (C) (200 nM each) and/or FV (400 nM, each) for 24 hours. WB membranes were probed with mouse monoclonal antibodies recognizing reduced FV, non-reduced uPA, and rabbit polyclonal Ab recognizing reduced VWF and β-actin (non-and HRP-conjugated, respectively). Nuclei were stained blue by DAPI. FV proteolysis in the presence of PLG alone is boxed in green and in the presence of PLG plus scuPA is in red or uPAT is in blue.

We then asked whether endocytosed uPAs would be activated by the presence of endocytosed native PLG and lead to degradation of endocytosed FV and/or endogenously expressed VWF. We found that the co-presence of endocytosed scuPA and PLG led to degradation of FV on Western blot, but not of endogenous VWF (Figure 5B). A similar study, but with added uPAT instead of scuPA, resulted in less FV degradation, and again, no cleavage of endogenous VWF (Figure 5C).

### In vivo studies of uPA-PLTs

We next examined *in vivo* in mice whether endocytosed uPA by *in vitro*-grown CD34^+^-MKs affects thrombopoiesis and subsequent platelet half-life and agonist-responsiveness. We utilized a model for studying the biology of released human platelets that we had previously described (Figure 6A). This model recapitulates the pulmonary release of platelets from megakaryocytes^44^ and where the released platelets closely resemble donor-derived platelets^25^. We found that both unmodified and uPA-exposed MK released a similar number of platelets per infused megakaryocyte, time to maximal platelet release, and platelet survival at 24 hours (Figure 6B). The uPA content per platelet appeared to remain stable over the study (Figure 6C) and uPA-MKs had similar responsiveness to human-specific, thrombin receptor activating peptide (TRAP) (Figure 6D).

**Figure 6.**
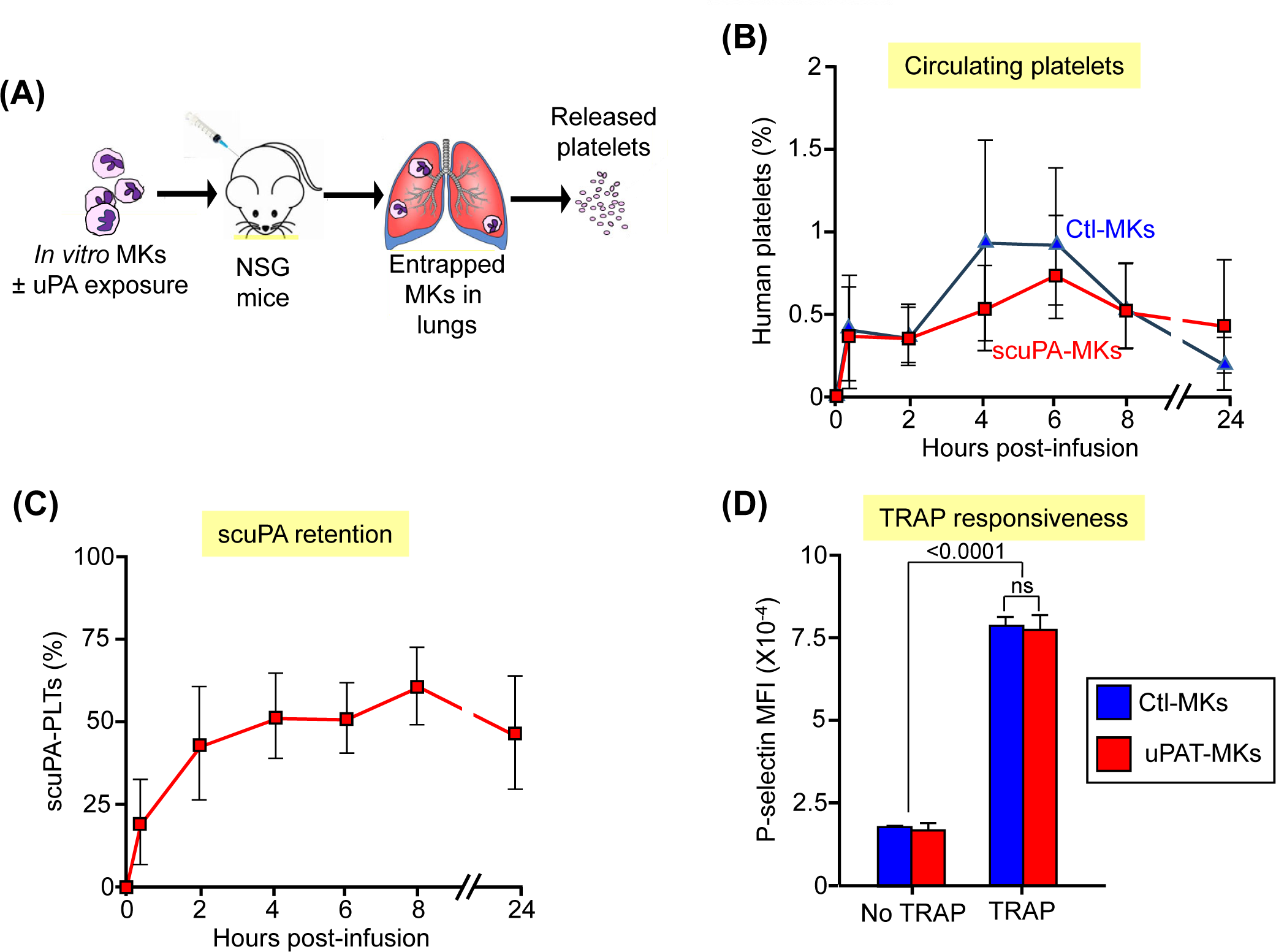
Release of human uPA-Plts in vivo after infusion of the intact and scuPA-loaded CD34+ MKs in NSG mice. **(A).** Schematic representation of intrapulmonary generation in NSG mice of human platelets from infused CD34^+^-MKs^64^ that had or had not been incubated to exogenous uPA. (**B**) 6×10^6^ d12-CD34^+^-intact MKs that had or had not been loaded with Alexa488-scuPA (400 nM) for 24 hours were injected into NSG mice. At each time point, peripheral blood sample was withdrawn and stained with APC-human CD41 and BUV395-mouse CD41 antibodies to measure circulating human platelets numbers relative to murine platelets. MFI was measured using flow cytometry. Y axis denotes % of human platelets (calculated from the total number of platelets measured in each blood sample) released at indicated times post-infusion of intact CD34^+^-Mks (blue) or -uPA-MKs (red). Mean ± 1 SD is shown. N = 8 (control Mks) and N = 6 (scuPA-MKs). (**C**). Same experiment as in (B), but scuPA retention was determined as the percent of human platelets that remained Alexa488-scuPA-positive at indicated time points. Mean ± 1 SD is shown. N = 6. (**D**) Studies of agonist responsiveness for the control MKs vs. uPAT-loaded MKs. Flow cytometry studies of the control MKs vs uPAT-loaded MKs at 8 and 24 hours after adding uPAT (400 nM) to the medium. P-selectin exposure after activation of the control and uPAT-loaded MKs the thrombin receptor activating peptide (TRAP) (50 μg/ml) was measured using FITC-anti-human CD41 and APC-anti-P-selectin monoclonal antibody. Mean ± 1 SD is shown. N = 3 per arm. * = P ≤ 0.0001 by one-way ANOVA comparing TRAP responsiveness of untreated MKs vs. uPAT-MKs.

*In vivo*, we also examined whether delivery of uPA via platelets would mediate thrombolysis in mice. To focus on uPA released by human platelets, we studied NSG mice and NSG transgenic mice that are homozygous for a mutant murine VWF that binds to the human glycoprotein Ib/IX, but not mouse. These NSG/VWF^R1326H^ mice exhibit a mild-to-moderate bleeding diathesis characterized by incomplete vessel occlusion and greater blood flow after photochemical carotid artery thrombosis was induced by rose bengal compared to NSG mice (Figure 7B, left-most two bars). After injection of human CD34^+^-MKs into NSG/VWF^R1326H^ mice, carotid artery occlusion improved to be similar to that seen in NSG mice. In contrast, injection of CD34^+^-MKs pre-loaded with either scuPA or uPAT prolonged the time to occlusion to nearly that seen with no human megakaryocyte infusion, consistent with endocytosed uPA in MKs injected in NSG/VWF^R1326H^ preventing nascent thrombus growth after induction of a vascular injury.

**Figure 7.**
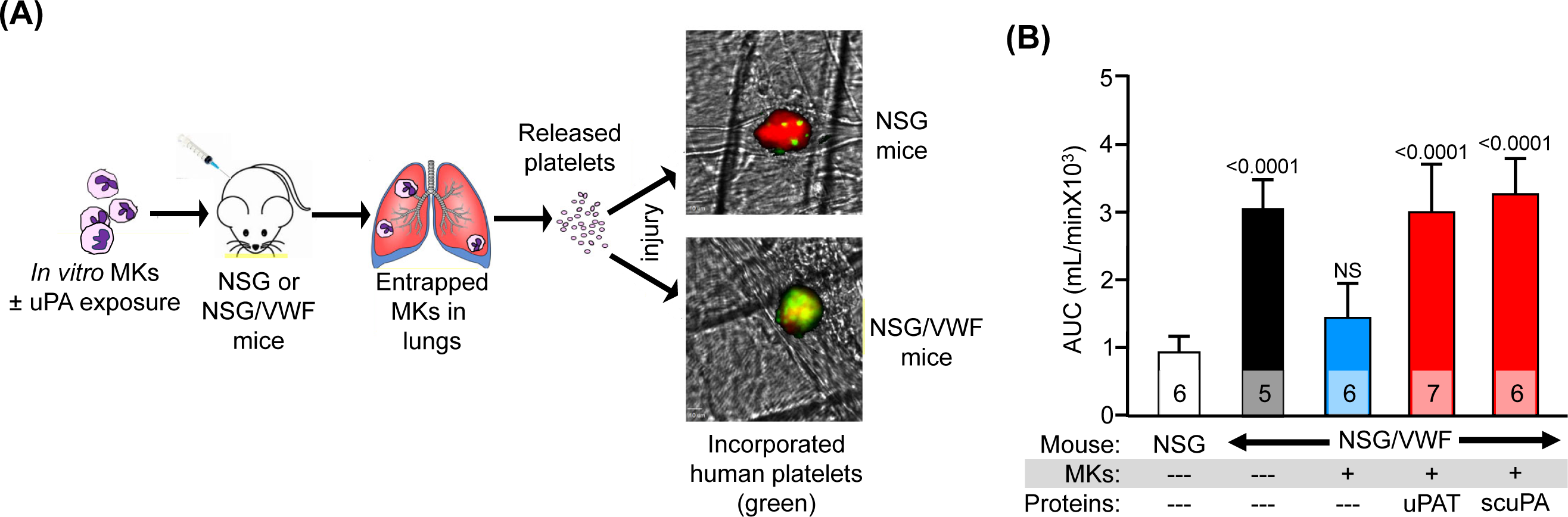
Anti-thrombotic effects of uPA-PLTs. (**A**) Schematic representation of the *in vivo* thrombosis model. d12-CD34^+^-MKs that had been loaded with scuPA or uPAT for 24 hour were injected in NSG or NSG/VWF mice. Infused MKs are trapped in the lung vasculature where they release platelets^64^. Photochemical intravascular injury is induced by injection of rose bengal and exposure to a 540 nm laser^31^. Human platelet released from CD34^+^-MKs (green) are incorporated into the nascent thrombus also containing murine platelets (red). (**B**) Area under the curve (AUC) of blood flow over the first 40 minutes post injury. The genotype of the recipient mice and the infusion of CD34^+^-MKs are shown in the X-axis as well whether the megakaryocytes had been incubated in uPA. The mean ± 1 SD and the number of independent experiments are shown in each bar. P values show outcomes in NSG/VWF studies compared with NSG control mouse (white bar) as determined by ordinary one-way ANOVA analysis.

## Discussion

For several years, we pursued a promising approach to targeted thrombolysis using platelets to deliver a fibrinolytic agent to sites of nascent thrombosis, avoiding lysis of mature thrombi. We called such platelets “anti-thrombotic thrombocytes”. Our prior *in vivo* studies in mice showed that infusing the uPA-containing platelets, isolated from transgenic mice that expressed scuPA during megakaryopoiesis into wildtype mice led to clot lysis without causing bleeding^14^; however, it was not clear how we could translate this genetic approach to clinical care and circumvent the excessive α-granule protein proteolysis by uPA-generated plasmin, seen in QPD^16–19^. The development of technologies to generate donor-independent *in vitro*-generated megakaryocytes^45,46^ and potentially platelets has led to the first description of infused *in vitro*-generated platelets in a patient^23^ and offered us a strategy to re-visit our “anti-thrombotic thrombocytes” approach. We envision that such generated platelets can be incubated with uPA prior to infusion to a patient to achieve targeted fibrinolysis at the sites of incipient thrombosis.

We tested two different uPAs: *i.* scuPA being a full-length uPA that retains plasmin activation, but is known to be able to be taken up by uPAR^47,48^ as well as other receptor systems on megakaryocytes that are retained in N-terminally truncated low-molecular weight uPA variants^49,50^, and *ii.* uPAT, a low-molecular weight uPA that lacks a plasmin activation site, but instead can be activated by thrombin^51^. Our reason to study uPAT was to limit proteolysis of uPA in the platelet α-granule pool that is seen in QPD platelets^16–19^. Another reason is that by requiring thrombin to be present would limit activation of uPAT to actively thrombosing sites where thrombin would be available as opposed to mature thrombus sites.

Surprisingly, scuPA and uPAT have similar endocytic characteristics into megakaryocytes. Both are targeted to the same granular pools in *in vitro*-differentiated megakaryocytes. Uptake of these uPAs appears to overlap with the uptake of the naturally endocytosed proteins, FV and fibrinogen, with considerable overlap in the granules in which they are stored. About half of FV endocytosis had been reported to occur via LRP1^36^, and we show the same for the uPAs. We also show that the uPAs are also taken up to a similar extent via αIIbβ3-binding. The LRP1 and αIIbβ3 pathways are not completely independent as the sum total of uptake via these two pathways never exceeds approximately half of the total uptake of the uPAs. Whether the remaining uptake by megakaryocytes of the uPAs, FV and fibrinogen involves a less-specific mechanism like pinocytosis needs to be examined^52,53^.

Our initial idea that motivated these *in vitro*-grown megakaryocyte studies that their growth in a specified cultured medium, lacking the exogenous proteins normally taken up from the marrow stroma, leads to “naked α-granules” appears to be supported. Our studies showed that the addition of FV and fibrinogen to the culture medium competes for uPA uptake and that the highest level of uPA uptake occurs when no competing protein is added to the culture medium (Figure 2). Whether uptake is limited by α-granule storage capacity or the availability of surface receptors and their limited capacity to recycle to the surface during megakaryopoiesis is untested. What is clear is that the capacity to endocytose uPAs is lost by released platelets, likely due to the fact that receptors, such as LRP1, are no longer present on the platelet surface (Figures 3A and S2). Thus, megakaryocytes can store functionally significant amounts of a protein normally not present in these cells and form modified platelets capable of performing a unique biological function distinct from a role in hemostasis.

Our studies also support that uPAT would be at less risk of proteolysis of α-granular proteins than scuPA. We show that the risk is greatest for other endogenous proteins and less so for endogenous proteins like VWF. This is distinct from what is seen in QPD and in our transgenic mice expressing murine urokinase where multiple exogenous and endocytosed proteins were shown to be proteolyzed^14,35,54^. We propose that PLG is initially endocytosed by megakaryocytes into an endocytosis α-granule compartment, but with time, these admixed with an endogenous α-granule compartment in the megakaryocytes and in the subsequent platelets. Our studies with added scuPA and PLG are too short-lasting to see the subsequent admixing of the two original pools, and so while endocytosed FV was clearly digested, VWF in the endogenous α-granule pool was protected, while with added uPAT and PLG endocytosed FV is more protected from degradation (Figures 5B and 5C, respectively).

Our murine *in vivo* data suggest that the exposure of developing megakaryocytes to uPA does not alter their capacity to release functional platelets post-infusion (Figure 6). Both scuPA-PLT and uPAT-PLT appear to be equally effective in preventing significant thrombus growth at a site of photochemical injury in a carotid artery model (Figure 7B). While in the presence of added PLG, scuPA-MKs may proteolyze α-granule proteins, in the absence of PLG addition, scuPA as well as uPAT would not be activated, leaving all α-granule proteins intact. Whether having a thrombin-activatable site in uPAT, rather than a plasmin-activatable site in scuPA, results in more targeted fibrinolysis to site of new thrombi and allows greater discrimination from mature thrombi needs to be tested.

Finally, besides targeted storage of uPAs in platelets, these strategy of delivery of a protein to a site of vascular injury has been proposed for FVIII in hemophilia A^55,56^ and ADAMTS13 in thrombotic thrombocytopenic purpura^57,58^. It may also be useful to deliver anti-angiogenic proteins in prevention of hematogenous cancer spread or anti-inflammatory proteins in a number of thromboinflammatory disorders. The further development of a practical system for *in vitro*-platelet preparation, we believe, is the major limitation. Once such a system is accepted clinically, its use for targeted delivery of therapeutics of a wide variety could be pursued. Many of these uses may require much smaller pool of modified platelets than needed for maintaining hemostasis and platelet counts.

In summary, *in vitro*-grown megakaryocytes in defined media can endocytose an ectopic protein such as the uPAs, scuPA and uPAT. The uptake of these uPAs are by pathways that overlap with those used by naturally endocytosed proteins, like FV and fibrinogen, and uPA uptake can be competed with the addition of these proteins. uPA uptake can be partially blocked by inhibiting uptake through LRP1 using FcRAP and uptake via αIIbβ3 using Abciximab. These two overlapping pathways only account for half of uPA uptake. The basis for the remaining uptake is unclear at the moment. The endocytosed uPAs are in a distinct α-granule pool from those containing endogenously expressed proteins like VWF. PLG is also endocytosed by megakaryocytes and its co-uptake with scuPA and FV leads to complete FV proteolysis, but its co-uptake with uPAT leads to limited FV uptake. In these studies, VWF is not proteolyzed consistent with their segregation to a distinct α-granule pool. *In vivo* studies show that thrombopoiesis and platelet biology is not affected by megakaryocyte incubation with these uPAs, and the resulting platelets can prevent nascent thrombi formation in mice. These studies suggest a new strategy for generating targeted therapy using *in vitro*-generated platelets that may be of benefit for selective fibrinolysis and serve as a model for potential other clinical uses.

## Supporting information

combined supplement file

## Acknowledgment

Support for this paper came from the following NIH grants: R35 HL150698 (MP), RO1HL141462 (VS), RO1HL159256 (DBC), P01 HL139420, Project 2 (RMC).

## Contributions to manuscript

MP along with VS provided overall leadership and experimental design and manuscript preparation. SZ prepared and characterized uPAs and performed the confocal and Western blot studies. He contributed to the data analysis and writing. HSA carried out the cell growth studies and the flow cytometric and *in vivo* studies. She contributed to the data analysis and writing. MAK, KB and RC contributed important ingredients for these studies. DBC made valuable insights to the studies and their interpretation as well as manuscript preparation.

