## Supplementary material for "Packaging of supplemented urokinase into naked alpha-granules of *in vitro*-grown megakaryocytes for targeted therapeutic delivery": combined supplement file

**SUPPLEMENTAL DATA**

### Packaging of uPA in *in-vitro*-grown CD34+ megakaryocytes as a novel strategy for platelet-mediated targeted delivery of fibrinolytics

Mortimer Poncz[1,2], Sergei Zaitsev[1], Hyunsook Ahn[1], M. Anna Kowalska[1], Khalil Bdeir[3], Rodney Camire[1,2], Douglas B. Cines[3], Victoria Stepanova[3]

Departments of [1]Pediatrics, The Children's Hospital of Philadelphia; and Departments of [2]Pediatrics and [3]Pathology and Laboratory Medicine and Medicine, University of Pennsylvania - Perelman School of Medicine, Philadelphia, PA 19104.

**SUPPLEMENT FIGURE LEGENDS.**

**Figure S1.** Dose-dependent (A) and time course (B) uptake of Alexa$^{488}$-uPAT (400 nM) uptake by CD34$^{+}$-Mks. Y axes denote mean fluorescence intensity (MFI) measured by flow cytometry. Mean ± 1 SD are shown for N=4 independent studies.

**Figure S2.** Flow cytometry analysis of endocytosed Alexa$^{488}$-uPAT (400 nM) by d10 CD34$^{+}$-MKs vs. isolated hPLTs. Y axes denote mean fluorescence intensity (MFI) measured by flow cytometry.

**Figure S3.** Confocal image of d11-CD34$^{+}$-MKs preincubated with uPAT (600 nM) for 24 hours starting at d10, fixed with 4% PFA in PBS, permeabilized with 0.1% Triton X-100 and stained with mouse anti-uPA MAbs (IMTEK) and anti-vWF Ab (Dako) followed by Alexa 488 goat anti-mouse Ab (green) and Alexa647 goat anti-rabbit Ab (red) and DAPI nuclear stain (blue). Scale Bar = 10 μm.

**Figure S4.** WB analysis of lysates prepared from d11-CD34$^{+}$-Mks with our without loading with enzymatically active PLG (20 μg/ml) on d10 for 18 hours. WB membranes were probed with mouse monoclonal Abs recognizing PLG (Santa Cruz Biotechnology) and HRP-conjugated anti-β-actin rabbit polyclonal Ab (CST).

**Major Resources Tables**

**Antibodies**

Summary of primary antibodies used for immunofluorescence (IF) and immunoblot analysis (WB)

| Type | Antigen | Source | Product #/Clone |
|---|---|---|---|
| **Rabbit** | | | |
| | vWF | Dako | A0082 |
| | mouse uPA | Meridian | K63679R |
| | b-actin HRP | CST | # 5125 |
| | LRP-1 | Novus Biologicals | SA0290 |
| | IFITM-3 | ProteinTech | CL594-11714 |
| | Total rabbit IgGs | Jackson ImmunoResearch | |
| **Mouse** | | | |
| | human uPA | IMTEK | |
| | PLG | SCBT | |
| | Factor V | Hematologic Technologies | Clone AHV-5146 |
| | P-selectin APC | BD Pharmingen | #550888 |
| | hCD41a APC | BD Pharmingen | #559777 |
| | hCD41a FITC | BD Pharmingen | # 555466 |
| | mCD41 BUV395 | BD Pharmingen | #565980 |
| | Total mouse IgGs | Jackson ImmunoResearch | # 015-000-003 |
| **goat** | | | |
| | Anti mouse HRP<br><br>Anti-rabbit HRP | Jackson ImmunoResearch<br>Jackson ImmunoResearch | |
| **recombinant** | | | |
| | Reo-Pro (Absiximab) | Centocor B.V. | CAS #: 143653-53-6 |

**Recombinant Proteins**

| Protein | Type | Conjugate | Source |
|---|---|---|---|
| scuPA | recombinant | None<br>Alexa488<br>Alexa555<br>Alexa568 | In-house |
| uPAT | recombinant | Alexa488 | In-house |
| Factor V | recombinant | None<br>Alexa568 | In-house |
| ncPLG | recombinant | Alexa647 | In-house |
| PLG | purified | None | Enzyme Research Labs |

| | | | |
|---|---|---|---|
| Fibrinogen | purified | Alexa488 | Hyphen/Aniara |
| FcRAP | recombinant | None<br>Alexa555 | In-house |

Fig. S1

### Dose dependence and time course of uPA-T uptake by CD34+ MKs

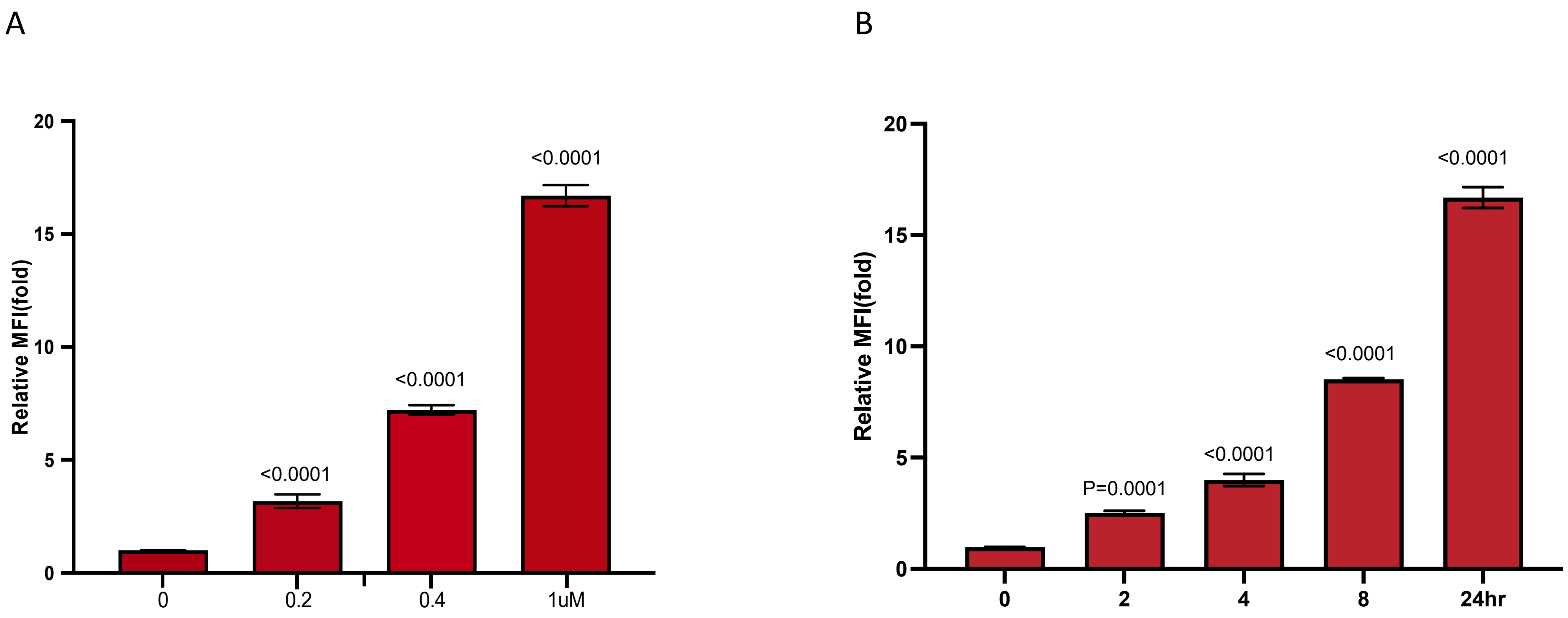

Fig. S2

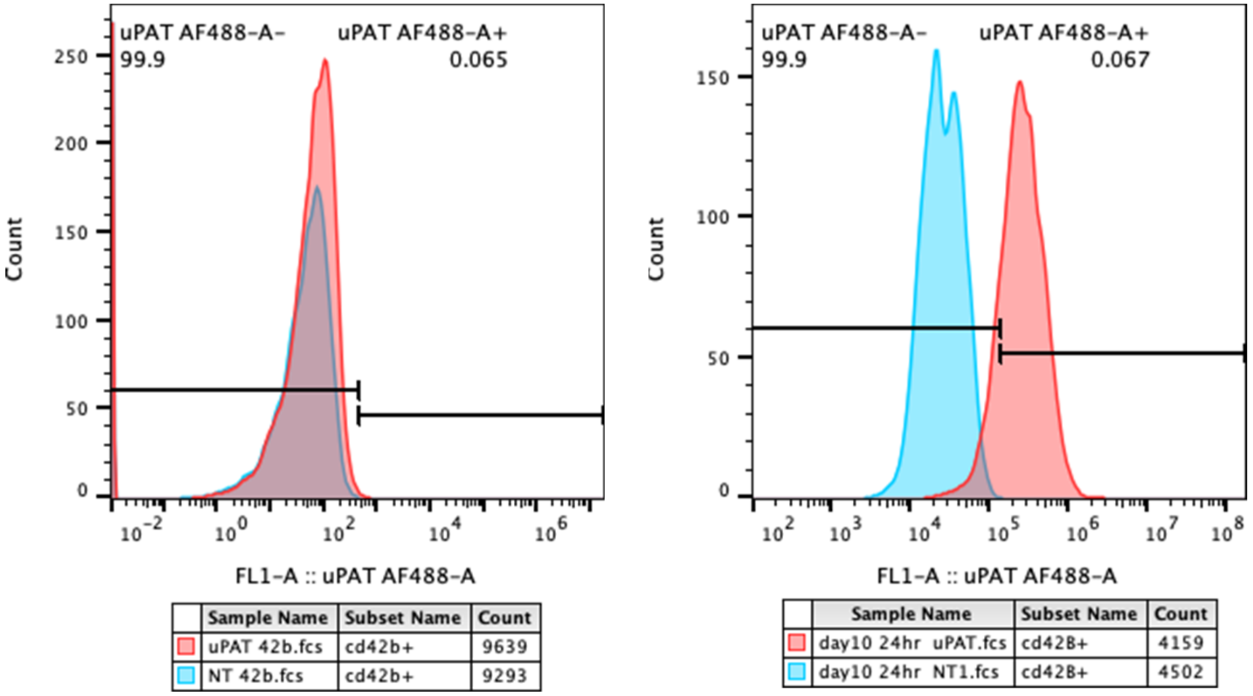
uPAT AF488-A-
99.9
uPAT AF488-A+
0.065
Count
FL1-A :: uPAT AF488-A
Sample Name | Subset Name | Count
uPAT 42b.fcs | cd42b+ | 9639
NT 42b.fcs | cd42b+ | 9293
uPAT AF488-A-
99.9
uPAT AF488-A+
0.067
Count
FL1-A :: uPAT AF488-A
Sample Name | Subset Name | Count
day10 24hr uPAT.fcs | cd42B+ | 4159
day10 24hr NT1.fcs | cd42B+ | 4502

Fig. S3

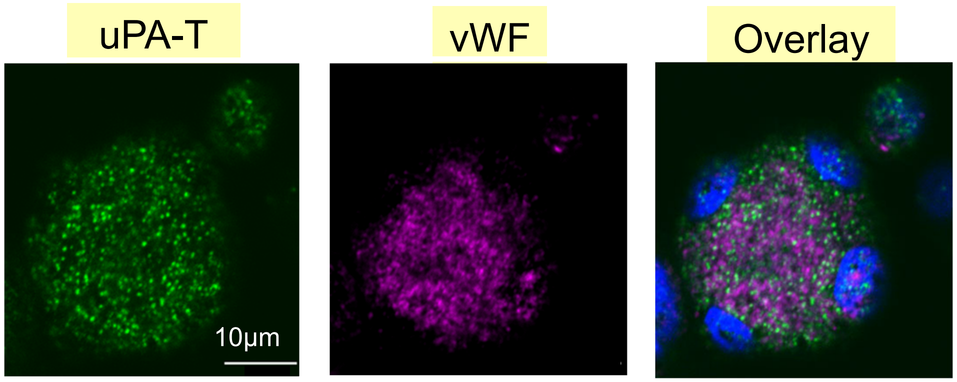

Fig. S4

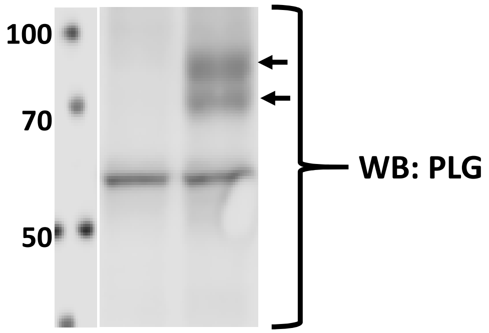
